# ERM proteins support perinuclear actin rim formation

**DOI:** 10.1101/2025.02.22.639617

**Authors:** Andrea Fracchia, Yuval Hadad, Dagmawit Babele, Gabi Gerlitz

**Affiliations:** Department of Molecular Biology, Faculty of Life Sciences and Ariel Center for Applied Cancer Research, Ariel University, Ariel 40700, Israel

**Author notes:** Correspondence to: Department of Molecular Biology, Faculty of Natural Sciences, Ariel University, Kiryat Hamada, Ariel 40700, Israel.

**Keywords:** nuclear envelope, Radixin, Moesin, Ezrin, Emerin, cell migration, LINC complex

## Abstract

The interaction of actin filaments with the nuclear envelope is essential for diverse cellular processes, including cell migration, nuclear positioning, and transcriptional control. The main studied mechanism that links F-actin to the nucleus is the Linker of Nucleoskeleton and Cytoskeleton (LINC) complex. Recently, the formation of a perinuclear actin rim has been identified in various cell types in response to external force or migration signals. This rim depends on the activation of the actin nucleator Inverted formin 2 (INF2) by calcium influx. However, it is not clear how the rim is coupled with the nuclear envelope. Here we show that the nuclear membrane protein Emerin, which has an actin-binding domain, is not required for the perinuclear actin rim formation. Interestingly, we found that the Ezrin-Radixin-Moesin (ERM) proteins, known to link actin filaments to the cell membrane, are also localized to the nuclear envelope in melanoma cells. Overexpression of ERM proteins increased the perinuclear actin rim levels, while knockdown of ERM proteins led to a reduction in the rim levels. Thus, the ERM proteins appear as part of the mechanism that links actin filaments to the nuclear envelope.

## Introduction

Interaction of the actin network with the nuclear envelope is crucial for cell migration, correct positioning of the nucleus in polarized cells, mechanotransduction, and transcriptional control (Davidson and Cadot, 2021). One of the most recently identified nuclear envelope-associated actin structures is the perinuclear actin rim, which is also termed Calcium-mediated Actin Reset (CaAR) (Shao et al., 2015b; Wales et al., 2016; Fracchia et al., 2020; Fracchia and Gerlitz, 2022; Jessop et al., 2024). The perinuclear actin rim is composed of actin filaments that engulf the nuclear envelope from its cytosolic side in both two-dimensional (2D) (Shao et al., 2015b; Wales et al., 2016; Fracchia et al., 2020) and 3D culture conditions (Fracchia et al., 2020). It was identified in various cells, including breast cancer cells, fibroblasts, and melanoma cells (Shao et al., 2015b; Wales et al., 2016; Fracchia et al., 2020; Jessop et al., 2024). In some cells, the perinuclear actin rim is formed transiently for 1-5 minutes as a reaction to an external force applied to the cell. The mechanical force leads to calcium ion influx that activates the actin nucleator Inverted formin 2 (INF2) (Shao et al., 2015b; Wales et al., 2016). INF2 is a member of the formin family that supports actin polymerization, which can localize to the endoplasmic reticulum (Labat-de-Hoz and Alonso, 2020). In mouse melanoma cells, the perinuclear actin rim is much more stable and formed for hours upon release of cells from contact inhibition (Fracchia et al., 2020).

The perinuclear actin rim was suggested to affect the migration capabilities of cells. On the one hand, enhancing perinuclear actin rim formation by the addition of ATP that mediates calcium influx accelerated the migration rate of breast cancer cells (Wales et al., 2016). On the other hand, interference with the perinuclear actin rim by expressing an active form of the actin severing protein Gelsolin at the nuclear envelope led to a reduction in the migration rate of melanoma cells (Fracchia et al., 2020). In search for additional factors that affect the perinuclear actin rim formation, it was found that lamin b at the nuclear side of the nuclear envelope interfered with perinuclear actin rim formation at the cytosolic side of the nuclear envelope (Fracchia et al., 2020). Perinuclear actin rim formation is independent of either the formin mDia2 (Shao et al., 2015a) or the Linker of Nucleoskeleton and Cytoskeleton (LINC) complex (Shao et al., 2015b; Fracchia et al., 2020). The LINC complex is a nuclear envelope complex composed of the inner nuclear membrane SUN domain proteins that bind the outer nuclear membrane KASH domain proteins. Inside the nucleus, the SUN domain proteins interact with the nuclear lamina and chromatin, while outside of the nucleus, the KASH domain proteins can bind various cytoskeleton elements, including actin filaments (Hieda, 2019; Fracchia and Gerlitz, 2022; King, 2023; McGillivary et al., 2023).

The accumulation of the perinuclear actin rim next to the nuclear envelope led us to hypothesize that an alternative mechanism to the LINC complex anchors the perinuclear actin rim to the nuclear envelope. Here, we looked for the involvement of factors that may help to connect the perinuclear actin rim to the nuclear envelope in mouse melanoma cells. We found that Emerin, a nuclear envelope protein that can bind F-actin to link it to the nuclear envelope in fibroblasts and keratinocytes (Le et al., 2016; Jin et al., 2023), is not necessary for the perinuclear actin rim formation. However, Ezrin-Radixin-Moesin (ERM) proteins that are known to be involved in connecting actin filaments to the plasma membrane (Ponuwei, 2016; Pelaseyed and Bretscher, 2018; Kawaguchi and Asano, 2022; Buenaventura et al., 2023) do localize to the nuclear envelope and support the formation of the perinuclear actin rim.

## Materials and methods

### Cell culture and transfection

Mouse melanoma B16-F10 cell line purchased from ATCC was grown as described previously (Maizels et al., 2017). Plasmids expressing GFP-fused human Ezrin, Radixin, and Moesin were a kind gift from Peter Vilmos. DNA plasmids transfection was done by the Nanojuice transfection kit (71900-3, Merck, Kenilworth, NJ, USA) and the jetOPTIMUS transfection reagent (101000025, Polyplus, Illkirch-Graffenstaden, France) following manufacturers’ instructions. Cells were incubated 24 hours before further analysis. For gene silencing, cells were transfected with siRNA (IDT, Coralville, IA, USA) with the INTERFERin reagent (101000028, Polyplus). Cells were incubated for 48 hours before further analysis. SiRNA used were mouse Emerin (mm.Ri.Emd.13.1), mouse Radixin (mm.Ri.Rdx.13.1), mouse Moesin (mm.Ri.Msn.13.2), mouse Ezrin (mm.Ri.Ezr.13.1) and negative control (51-01-14-04). SiRNA transfection efficiency was >90%, as verified by transfection of Cy3 Transfection Control DsiRNA (51-01-03-06).

### Immunostaining

Cells plated on Fibronectin (03-090-1-05, Biological Industries, Beit-Haemek, Israel) coated cover-glasses with or without a scratch were fixed by 2% paraformaldehyde for 10 min at room temperature. Antibodies used were rabbit anti-Emerin (sc-15378, Santa Cruz Biotechnology, Dallas, TX, USA) diluted 1:150, mouse anti-Ezrin (sc-58758, Santa Cruz Biotechnology, Heidelberg, Germany) diluted 1:200, goat anti-GFP (ab5450, Abcam, Cambridge, UK) diluted 1:400, goat anti-Lamin B (6216, Santa Cruz, Biotechnology, Dallas, TX, USA) diluted 1:150, mouse anti-Moesin (CST3150, Cell Signaling Technologies, Danvers, MA, USA) diluted 1:300 and rabbit anti-Radixin (ab52495, Abcam, Cambridge, UK) diluted 1:100. Actin filaments were labeled by DyLight™ 554 Phalloidin (13054, Cell Signaling Technologies, Danvers, MA, USA) diluted 1:400. DNA was stained with Hoechst 33342 (B2261, Sigma-Aldrich, Rehovot, Israel). Images were collected using an Olympus IX81 fluorescence microscope with a coolSNAP HQ2 CCD camera (Photometrics, Tuscon, AZ, USA) or a Prime BSI Express camera (Teledyne Photometrics, Tucson, AZ, USA). The ImageJ/Fiji software (National Institute of Health, Bethesda, USA) was used to measure Phalloidin mean intensities at the nuclear envelopes, as marked by Lamin B immunostaining. Images were assembled with Photoshop (Adobe, San Jose, CA, USA).

### Protein lysate preparation and Western blot analysis

Cells were washed in PBS, scraped, precipitated by centrifugation at 500 x *g* for 5 min at 4°C and sonicated in 2x SDS sample buffer (100 mM Tris pH 6.8, 10 % glycerol, 2 % SDS, 0.1 M DTT, and bromophenol blue) supplemented with protease inhibitor cocktail (539134, Merck, Kenilworth, NJ, USA). Samples were then heated at 95 ºC for 10 minutes and kept at - 20 ºC until use. Protein extracts were separated in SDS-PAGE and analyzed by Western blot analysis using the following antibodies: rabbit anti-CTCF (3418, Cell Signaling Technology, Danvers, MA, USA) diluted 1:1,000, rabbit anti-Emerin (sc-15378, Santa Cruz Biotechnology, Dallas, TX, USA) diluted 1:5,000, mouse anti-Ezrin (sc-58758, Santa Cruz Biotechnology, Heidelberg, Germany) diluted 1:1,000, rabbit anti-histone H3 (05-928, EMD Millipore, Temecula, CA, USA), mouse anti-Moesin (CST3150, Cell Signaling Technologies, Danvers, MA, USA) diluted 1:1,00 and rabbit anti-Radixin (ab52495, Abcam, Cambridge, UK) diluted 1:1,000. Images were assembled with Photoshop (Adobe, San Jose, CA, USA).

## Results

Perinuclear actin rim was found to be connected to the nucleus in a LINC complex independent manner in mouse fibroblasts and melanoma cells (B16-F10 cells) (Shao et al., 2015b; Fracchia et al., 2020). The nuclear envelope protein Emerin was shown to bind actin (Lattanzi et al., 2003; Holaska et al., 2004) and to support perinuclear actin accumulation in response to mechanical strain (Le et al., 2016; Jin et al., 2023). To test the possibility that Emerin is also essential for the perinuclear actin rim in B16-F10 cells, we first looked for its localization in these cells upon release from contact inhibition in the wound healing assay. As expected, we could detect Emerin mainly at the nuclear periphery and, to some extent, in the cytoplasm in a pattern that resembles the nuclear envelope and the endoplasmic reticulum (ER), respectively. This pattern of localization was reported before for Emerin (Salpingidou et al., 2007; Le et al., 2016). The knockdown of Emerin (Sup. Fig. 1) did not affect the intensity of the perinuclear actin rims in a significant manner in B16-F10 cells (Fig. 1). Suggesting that another factor links the perinuclear actin rim to the nucleus.

**Figure 1.**
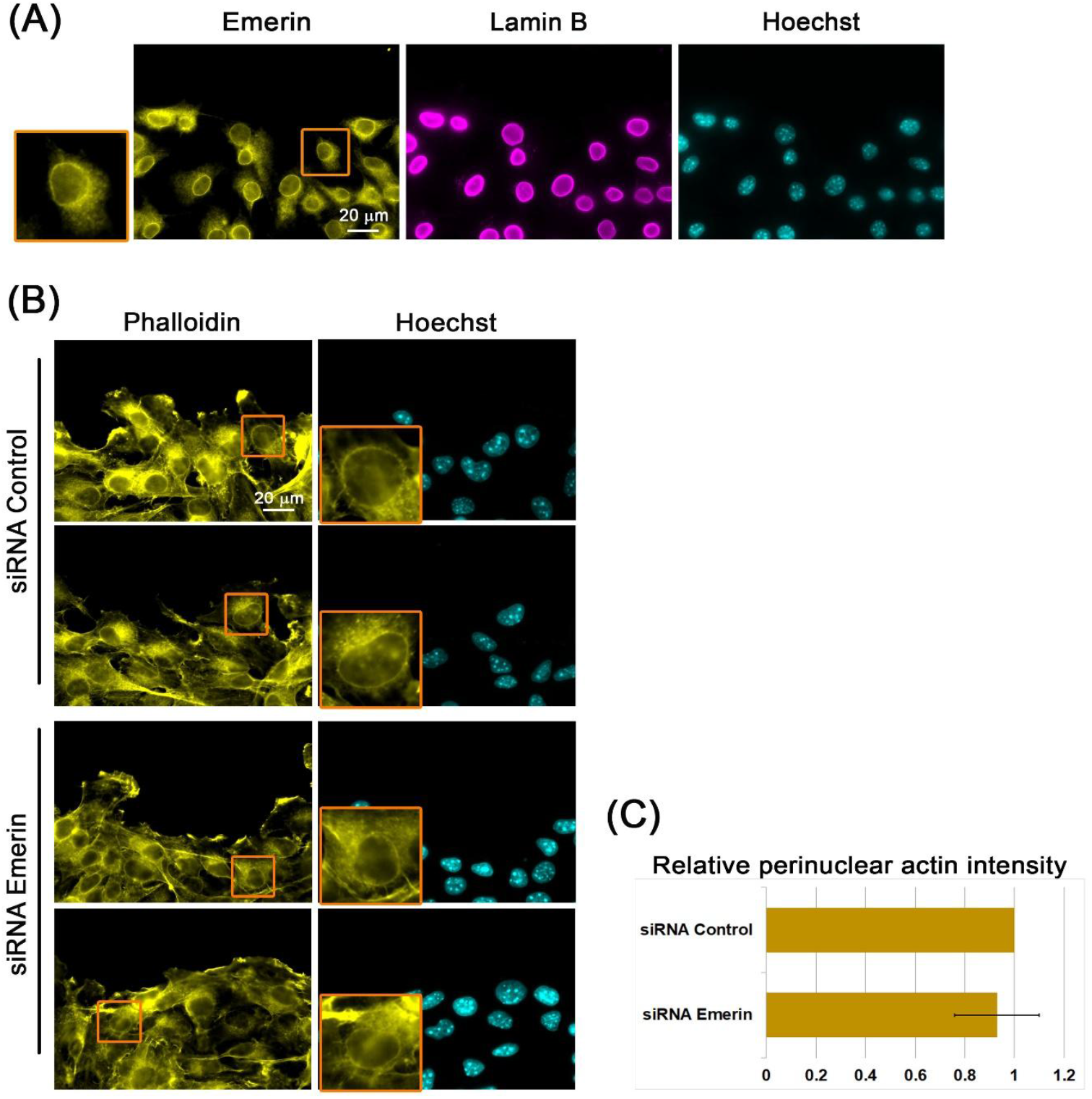
Emerin is dispensable for perinuclear actin rim formation. (**A**) Emerin in B16-F10 cells. Confluent B16-F10 cells induced to migrate in the wound healing assay for 3 h immunostained for Emerin, Lamin B, and DNA (Hoechst). The edge of the scratch is in the top region of each micrograph. Scale bar: 20 µm. The nucleus in the orange rectangle is magnified on the left side. (**B**) Actin perinuclear rim in Emerin KD cells. Confluent B16–F10 cells treated with either control or Emerin siRNA were induced to migrate in the wound healing assay for 3 h and stained for filamentous actin (Phalloidin) and DNA (Hoechst). The edge of the scratch is in the top region of each micrograph. The nuclei in the orange rectangles are magnified. Scale bar: 20 µm. (**C**) Quantification of the actin perinuclear rim in siRNA Control vs. siRNA Emerin treated B16–F10 cells. For quantification, in each experiment, 40–50 cells of each condition were measured for the Phalloidin signal at the nuclear periphery. The mean intensity was calculated and normalized to control cells. The sample difference was not statistically significant based on the Student’s *t*-test.

The ERM proteins are well-established factors involved in connecting actin filaments to the plasma membrane (Ponuwei, 2016; Pelaseyed and Bretscher, 2018; Kawaguchi and Asano, 2022; Buenaventura et al., 2023). To look for their possible involvement in perinuclear actin rim formation, we first checked for their intracellular localization. Notably, we could detect perinuclear accumulation of ERM proteins (Fig. 2).

**Figure 2.**
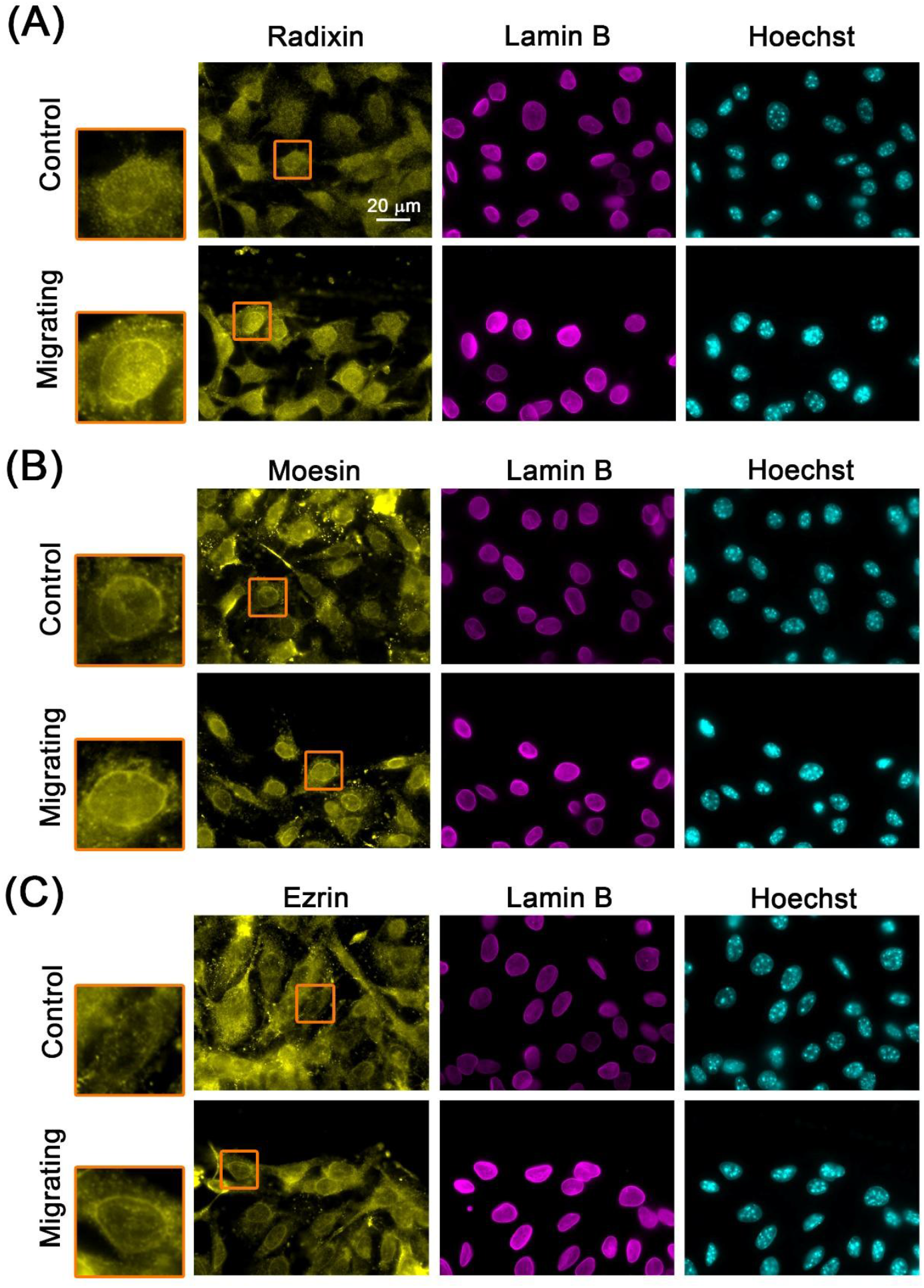
Perinuclear localization of ERM proteins. Confluent B16–F10 cells induced to migrate in the wound healing assay for 3 h immunostained for Radixin (A), Moesin (B), and Ezrin (C) along with Lamin B. DNA is stained with Hoechst. The edge of the scratch is in the top region of each micrograph. The nuclei in the orange rectangles are magnified. The merged images show the merged signals of ERM proteins and Lamin B. Scale bar: 20 µm.

This localization was found in both non-migrating and migrating cells. To check if ERM proteins can affect the perinuclear actin rim formation, we analyzed the actin rim intensity following overexpression of ERM proteins. As shown in Fig. 3, the over-expressed proteins were partially localized to the nuclear periphery. Notably, the intensity of the perinuclear actin rim increased significantly by 21% and 40% upon overexpression of GFP-fused Radixin and Moesin, respectively. This result suggests that ERM proteins can support the perinuclear actin rim.

**Figure 3.**
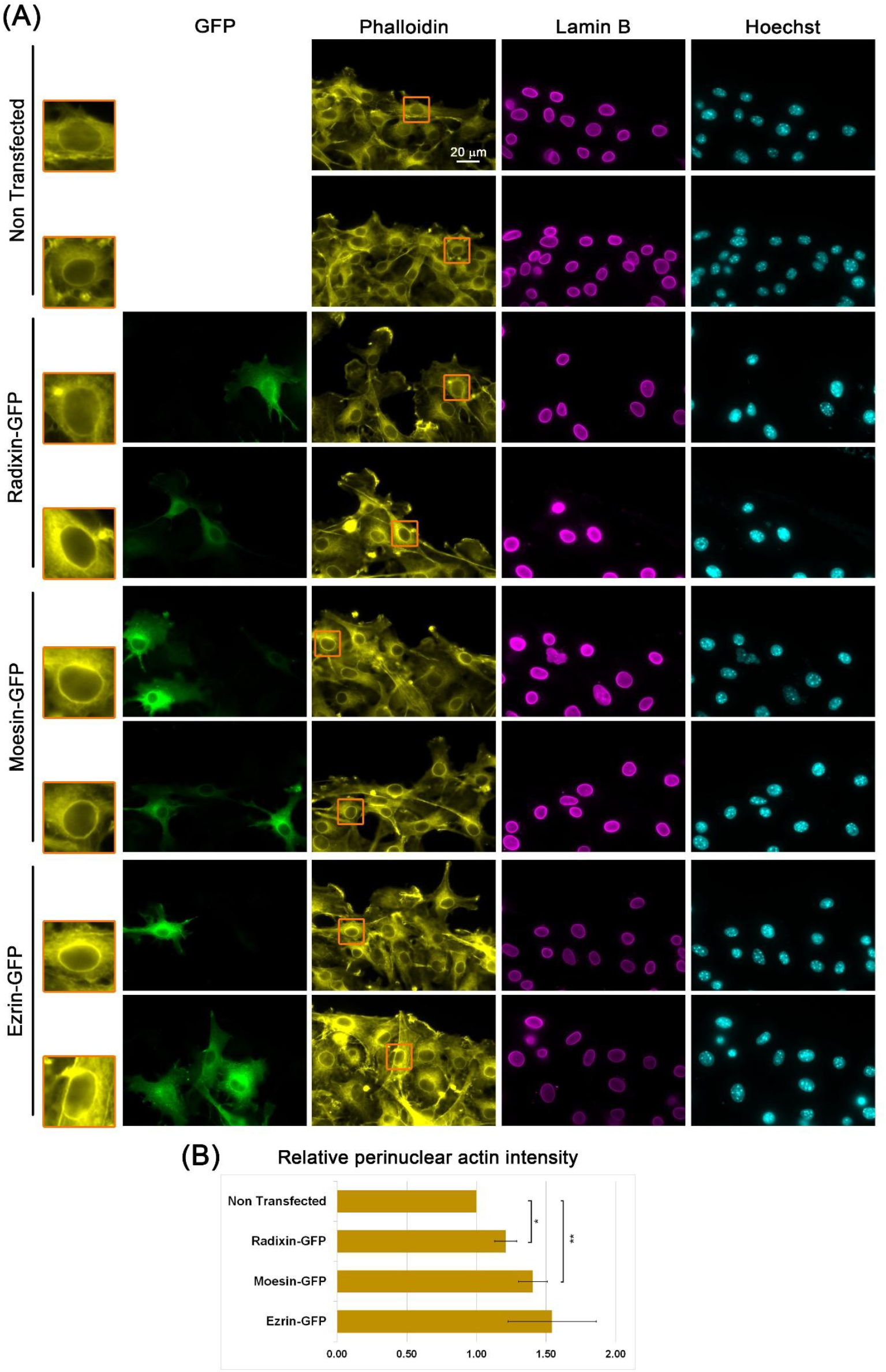
Overexpression of ERM proteins increases the intensity of the perinuclear actin rim. **(A)**Perinuclear actin rim upon overexpression of ERM proteins. Confluent B16-F10 over-expressing GFP-fused ERM proteins induced to migrate in the wound healing assay for 3 h stained for filamentous actin (Phalloidin), nuclear envelope (Lamin B) and DNA (Hoechst). DNA is stained with Hoechst. The edge of the scratch is in the top region of each micrograph. The nuclei in the orange rectangles are magnified on the left side. Scale bar: 20 µm. (**B**) Quantification of the actin perinuclear rim in ERM overexpressing cells vs. control cells. For quantification, in each experiment, 20–30 cells of each condition were measured for the Phalloidin signal at the nuclear periphery. The mean intensity was calculated and normalized to control cells. The average mean intensity in three independent experiments ± s.e. is presented. Statistical significance was evaluated by the Student’s *t*-test, \**P*<0.05, \*\**P*<0.01.

To verify this observation, we looked for the effect of ERM proteins knockdown (Sup. Fig. 2) on the formation of the perinuclear actin rim. We realized that the release of contact inhibition in the wound healing assay leads to perinuclear actin rim formation that lasts as long as the contact inhibition has not been restored. Therefore, at this point, we analyzed sub-confluent B16-F10 cells. As seen in Fig. 4, the intensity of the perinuclear actin rim was reduced by 25% and 50% following the knockdown of Moesin and Ezrin, respectively. These results support the hypothesis that ERM proteins are part of the molecular mechanism that generates the perinuclear actin rim.

**Figure 4.**
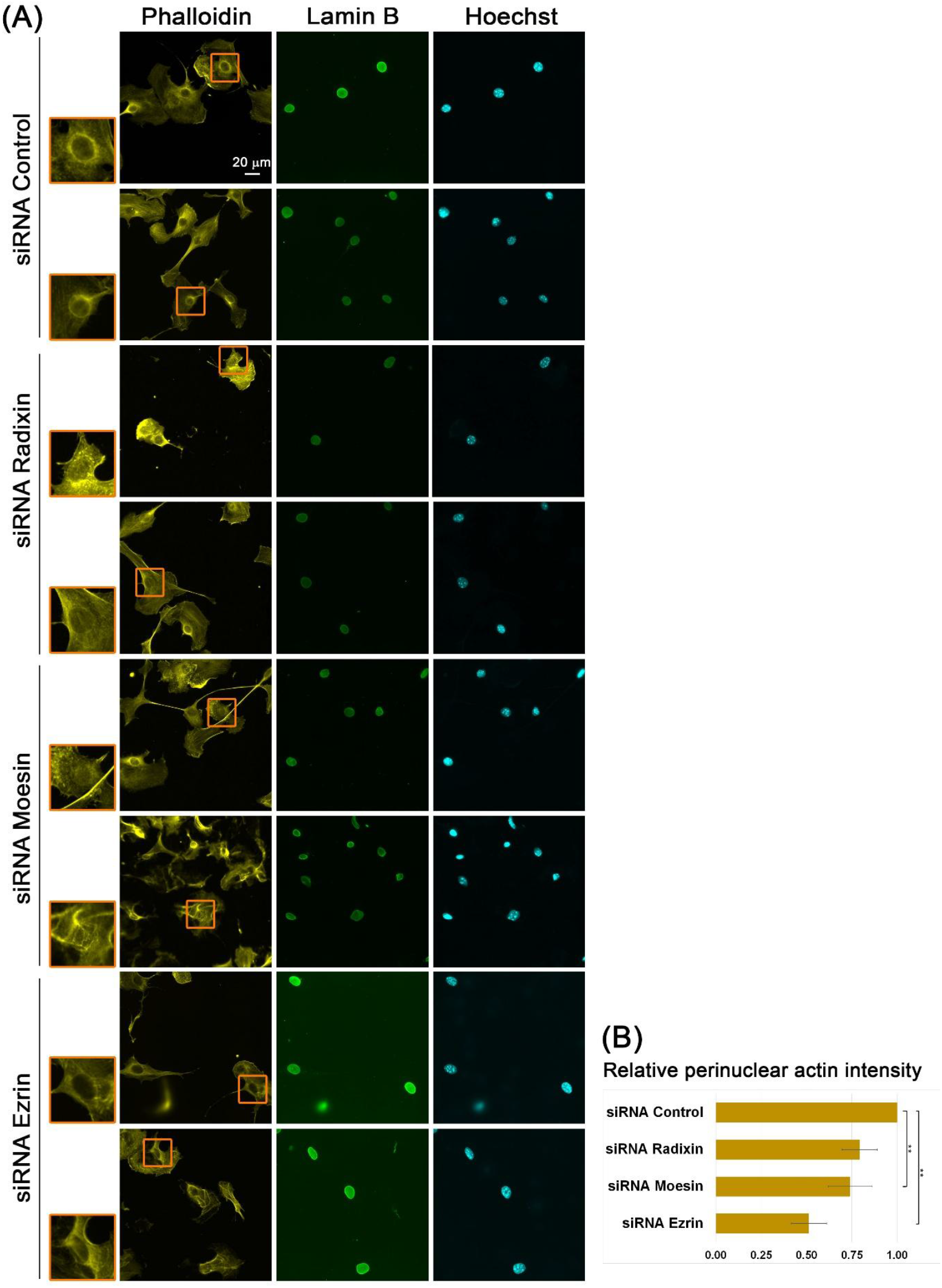
ERM proteins support the formation of a perinuclear actin rim. (**A**) Actin perinuclear rim in after KD of ERM proteins. Sub-confluent B16–F10 cells treated with either control, Radixin, Moesin, or Ezrin siRNA stained for filamentous actin (Phalloidin), nuclear envelope (Lamin B), and DNA (Hoechst). The nuclei in the orange rectangles are magnified on the left side. Scale bar: 20 µm. (**B**) Quantification of the actin perinuclear rim in siRNA Control vs. siRNA ERM proteins treated B16–F10 cells. For quantification, in each experiment, 20-30 cells of each transfection were measured for the Phalloidin signal at the nuclear periphery. The mean intensity was calculated and normalized to control cells. The average mean intensity in three independent experiments ± s.e. is presented. Statistical significance was evaluated by the Student’s *t*-test, \*\**P*<0.01.

## Discussion

The perinuclear actin rim has been proposed to affect both cytoplasmic processes, such as cell migration (Wales et al., 2016; Fracchia et al., 2020) and nuclear processes such as transcription (Wales et al., 2016). In search for the mechanism that links the perinuclear actin rim to the nuclear envelope the LINC complex was evaluated. However, interference with LINC complex function by overexpression of the KASH domain of either Nesprin 1 or Nesprin 2 that has a dominant negative effect did not affect the formation of the actin perinuclear rim (Shao et al., 2015b; Fracchia et al., 2020). Here, we looked for the possible involvement of Emerin and ERM proteins. Emerin can localize to the outer nuclear envelope, where it was shown to bind actin filaments (Le et al., 2016), however knockdown of Emerin did not affect the perinuclear actin rim (Fig. 1). ERM proteins can be found in the nucleus (Borkúti et al., 2024; Kovács et al., 2024) and Ezrin and Moesin were also identified in nuclear envelope fraction by a mass spectrometry analysis of a purified nuclear envelope from human leukocytes (Korfali et al., 2010). In our immunostainings, ERM proteins were partially localized to the nuclear periphery in a similar pattern to the lamin B immunostaining (Fig. 2). This localization suggests that ERM proteins can be associated not only with the plasma membrane but also with the nuclear membrane. Plasma membrane binding of ERM proteins is dependent on their FERM domain that interacts with phosphatidylinositol 4,5-bisphosphate [PI(4,5)P_2_] and membranal proteins (Pelaseyed and Bretscher, 2018; Kawaguchi and Asano, 2022; Buenaventura et al., 2023). Notably, PI(4,5)P_2_ was found not only in the plasma membrane but also in the nuclear membrane (Smith and Wells, 1983, 1984b, 1984a; Mazzotti et al., 1995; Watt et al., 2002; Fiume et al., 2019). Thus, PI(4,5)P_2_ in the nuclear membrane could recruit ERM proteins to engulf the nucleus. An additional activator of Ezrin is S100 calcium-binding protein P (S100P), which can bind and activate Ezrin in a calcium-dependent manner (Koltzscher et al., 2003; Austermann et al., 2008). It may be relevant since calcium inflex activates the formation of the perinuclear actin rim (Shao et al., 2015b; Wales et al., 2016). Although S100P has been mostly studied in relation to the plasma membrane, it can be found in the cytoplasm at the nuclear periphery and in the nucleus (Sakaguchi et al., 2000; Elevated nuclear S100P expression is associated with poor survival in early breast cancer patients, 2013).

Next, we evaluated the effect of altered levels of ERM proteins on the perinuclear actin rim. We found that overexpression of ERM proteins led to an increase in the perinuclear actin rim levels (Fig. 3). In support of this observation, the knockdown of ERM proteins led to reduced levels of the perinuclear actin rim (Fig. 4). In each intervention, two out of the three ERM proteins were found to affect significantly the perinuclear actin rim levels. These results suggest a redundancy among these proteins regarding perinuclear actin organization.

Taking together our data with other results, we suggest a model in which an increase in calcium ions at the nuclear periphery leads to activation of INF2 and polymerization of actin filaments that are held by nuclear membrane-localized ERM proteins to form a rim around the nucleus. The formed rim directly affects nuclear mechanical properties and indirectly affects transcription. The perinuclear actin rim formation was suggested to induce the release of serum response factor (SRF) co-factor MRTF-A from globular actin. This step exposes the nuclear localization signal (NLS) of MRTF-A, leading to its nuclear import and activation of target genes (Wales et al., 2016).

Here, we identify a possible new mechanism for linking actin filaments to the nuclear membrane by the ERM proteins. Our previous observation that lamin B at the inner side of the nuclear envelope can negatively affect the perinuclear actin rim formation (Fracchia et al., 2020) suggests the existence of a physical linkage between the ERM-linked actin filaments at the outer side of the nuclear envelope and the lamina at the inner side of the nuclear envelope that is still waiting to be discovered.

## Acknowledgments

We thank Peter Vilmos (Biological Research Center of the Hungarian Academy of Sciences, Szeged, Hungary) for providing plasmids.

## Authors Contributions

Conceptualization, A.F., Y.H., D.B. and G.G.; Investigation, A.F., Y.H. and D.B.; Writing – Original Draft Preparation, G.G.; Writing – Review & Editing, A.F., Y.H. and D.B.; Visualization, A.F., Y.H., D.B. and G.G.; Supervision, G.G.; Project Administration, G.G.; Funding Acquisition, G.G.

