## Supplemental figures for "ERM proteins support perinuclear actin rim formation"

### Supplementary Figures

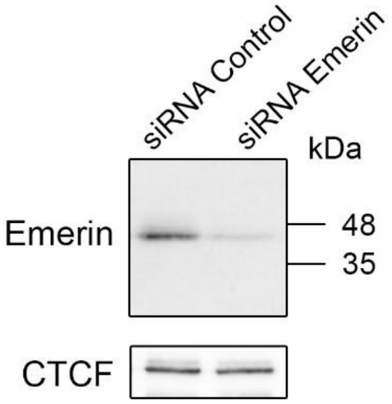


**Supplementary Figure 1. Emerin knockdown.** Western blot analysis of Emerin in control and Emerin siRNA-treated B16-F10 cells. CTCF was used as a loading control.

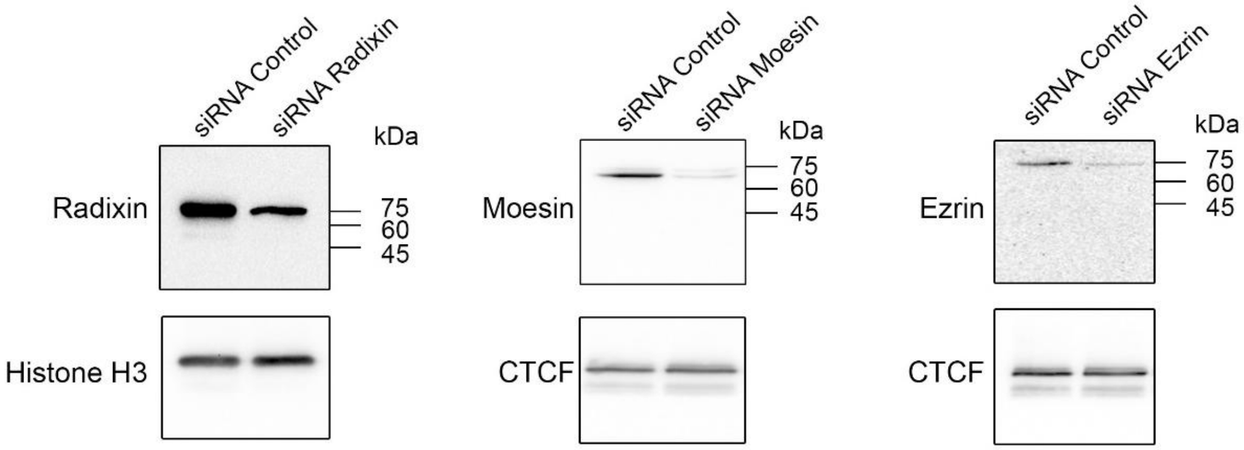


**Supplementary Figure 2. Knockdown of ERM proteins.** Western blot analysis of ERM proteins in control and Radixin/ Moesin/ Ezrin siRNA-treated B16-F10 cells. Histone H3 or CTCF was used as a loading control.
